# Durability of mRNA-1273-induced antibodies against SARS-CoV-2 variants

**DOI:** 10.1101/2021.05.13.444010

**Authors:** Amarendra Pegu, Sarah O’Connell, Stephen D Schmidt, Sijy O’Dell, Chloe A. Talana, Lilin Lai, Jim Albert, Evan Anderson, Hamilton Bennett, Kizzmekia S. Corbett, Britta Flach, Lisa Jackson, Brett Leav, Julie E. Ledgerwood, Catherine J. Luke, Mat Makowski, Paul C. Roberts, Mario Roederer, Paulina A. Rebolledo, Christina A. Rostad, Nadine G. Rouphael, Wei Shi, Lingshu Wang, Alicia T. Widge, Eun Sung Yang, the mRNA-1273 Study Group, John H. Beigel, Barney S. Graham, John R Mascola, Mehul S. Suthar, Adrian McDermott, Nicole A. Doria-Rose

**Affiliations:** Vaccine Research Center, National Institute of Allergy and Infectious Diseases, National Institutes of Health; Bethesda MD, USA; Department of Medicine, Center for Childhood Infections and Vaccines (CCIV) of Children’s Healthcare of Atlanta, Emory Vaccine Center, and Emory University Department of Pediatrics, Emory University School of Medicine; Atlanta, GA, USA; Emmes Company; Rockville, MD, USA; Moderna, Inc.; Cambridge, MA, USA; Division of Microbiology and Infectious Diseases, National Institute of Allergy and Infectious Diseases, National Institutes of Health; Bethesda, MD, USA; Kaiser Permanente Washington Health Research Institute; Seattle, WA, USA; Hope Clinic, Department of Medicine, Emory University School of Medicine; Decatur, GA, USA

## Abstract

SARS-CoV-2 mutations may diminish vaccine-induced protective immune responses, and the durability of such responses has not been previously reported. Here, we present a comprehensive assessment of the impact of variants B.1.1.7, B.1.351, P.1, B.1.429, and B.1.526 on binding, neutralizing, and ACE2-blocking antibodies elicited by the vaccine mRNA-1273 over seven months. Cross-reactive neutralizing responses were rare after a single dose of mRNA-1273. At the peak of response to the second dose, all subjects had robust responses to all variants. Binding and functional antibodies against variants persisted in most subjects, albeit at low levels, for 6 months after the primary series of mRNA-1273. Across all assays, B.1.351 had the greatest impact on antibody recognition, and B.1.1.7 the least. These data complement ongoing studies of clinical protection to inform the potential need for additional boost vaccinations.

**One-Sentence Summary:** Most mRNA-1273 vaccinated individuals maintained binding and functional antibodies against SARS-CoV-2 variants for 6 months.

## Main Text

SARS-CoV-2, the virus that causes COVID-19, has infected over 150 million people and resulted in over 3 million deaths globally as of early May 2021 (*1*). The combination of RNA virus mutability and replication in a very large number of individuals is conducive to the emergence of variant viruses with improved replication capacity and transmissibility, as well as immunological escape. Indeed, the B.1 variant (spike mutation containing D614G) replaced the initial circulating (prototypic virus Wuhan-Hu-1, also called WA1) strain by the summer of 2020 (*2*). Subsequently, other variants have emerged that may have increased transmissibility; the first variants that came to global attention were B.1.1.7 (also called 20I/501Y.V1), then B.1.351 (also called 20H/501Y.V2), which were in first identified in the United Kingdom and South Africa, respectively. Known as Variants of Concern, they bear 8 or 9 mutations in spike compared to the original Wuhan-Hu-1 strain, and share the N501Y mutation in spike. Notably, B.1.351 has a cluster of 3 mutations, K417N-E484K N501Y in the receptor-binding domain (RBD) that is associated with resistance to neutralization by monoclonal and polyclonal antibodies (*3, 4*). B.1.1.7 has been shown to be 2-3 fold less sensitive to sera from convalescent patients infected with WA1 or D614G, as well as recipients of vaccines derived from WA1, which is the parent sequence for all currently authorized vaccines (*4-8*). In contrast, B.1.351 is partially resistant to neutralization, with 6-15 fold less neutralization activity for sera from individuals vaccinated with WA1-based vaccines (*3, 6, 7, 9-11*). More recently, Variants of Concern P.1 (first identified in Brazil) and B.1.429 (also called Cal20, first identified in California) and Variant of Interest B.1.526 (first identified in New York) have been shown to have modest levels of resistance to convalescent or vaccine sera (*10, 12-15*). These prior studies have evaluated vaccine sera from vaccinated individuals only at timepoints soon after the first or second dose of various vaccines. Likewise, while clinical studies have reported efficacy and effectiveness against the B.1.1.7 and B.1.351 variants, these have been in the first several months following vaccination (*16, 17*)). Although such data provide critical insights into the performance of the vaccines against these viral variants, they have not addressed the durability of cross-reactive binding and functional antibodies.

Here we investigate the impact of these SARS-CoV-2 variants on recognition by sera from subjects who received two 100 mcg doses of the SARS-CoV-2 vaccine mRNA-1273 and were followed over 7 months after the first dose. We used three methodologies to measure functional characteristics – an ACE2 blocking assay, and SARS-CoV-2 pseudovirus and live-virus neutralization assays – and two methodologies to measure antibody binding to full length SARS-CoV-2 spike and soluble spike proteins, to comprehensively assess vaccine-elicited humoral immunity over time.

## Results

We previously described the binding and neutralization activity against the original SARS-CoV-2 Spike, herein referred to as WA1, longitudinally over 7 months from the first vaccination in volunteers from the Phase 1 trial of the mRNA-1273 vaccine (*18-21*). mRNA-1273 encodes the full-length stabilized spike protein of WA1 (also called Wuhan-Hu-1) sequence and was administered as a two-dose series 28 days apart. In the current study, we expanded the serologic evaluation to include additional assay formats and multiple variant forms of spike. We tested sera from a random sample of 8 volunteers in each of three age groups: 18-55, 55-70, and 71+ years of age, all of whom received the 100 mcg dose of vaccine and had samples available from four timepoints: 4 weeks after the first dose, and two weeks, 3 months, and 6 months after the second dose (Days 29, 43, 119, and 209 after the first dose, respectively).

We initially assessed the serum activity using a lentivirus-based pseudovirus neutralization assay to measure serum activity against D614G, B.1.1.7, and B.1.351 (Figure 1). Neutralization of D614G was slightly higher than previously reported for WA1 (*18, 20*), with all subjects maintaining activity up to Day 209. B.1.1.7 showed a similar pattern. B.1.351, however, was significantly less sensitive to serum neutralization, with serum titers in many subjects declining below the limit of detection at the later timepoints (Figure 1).

**Figure 1.**
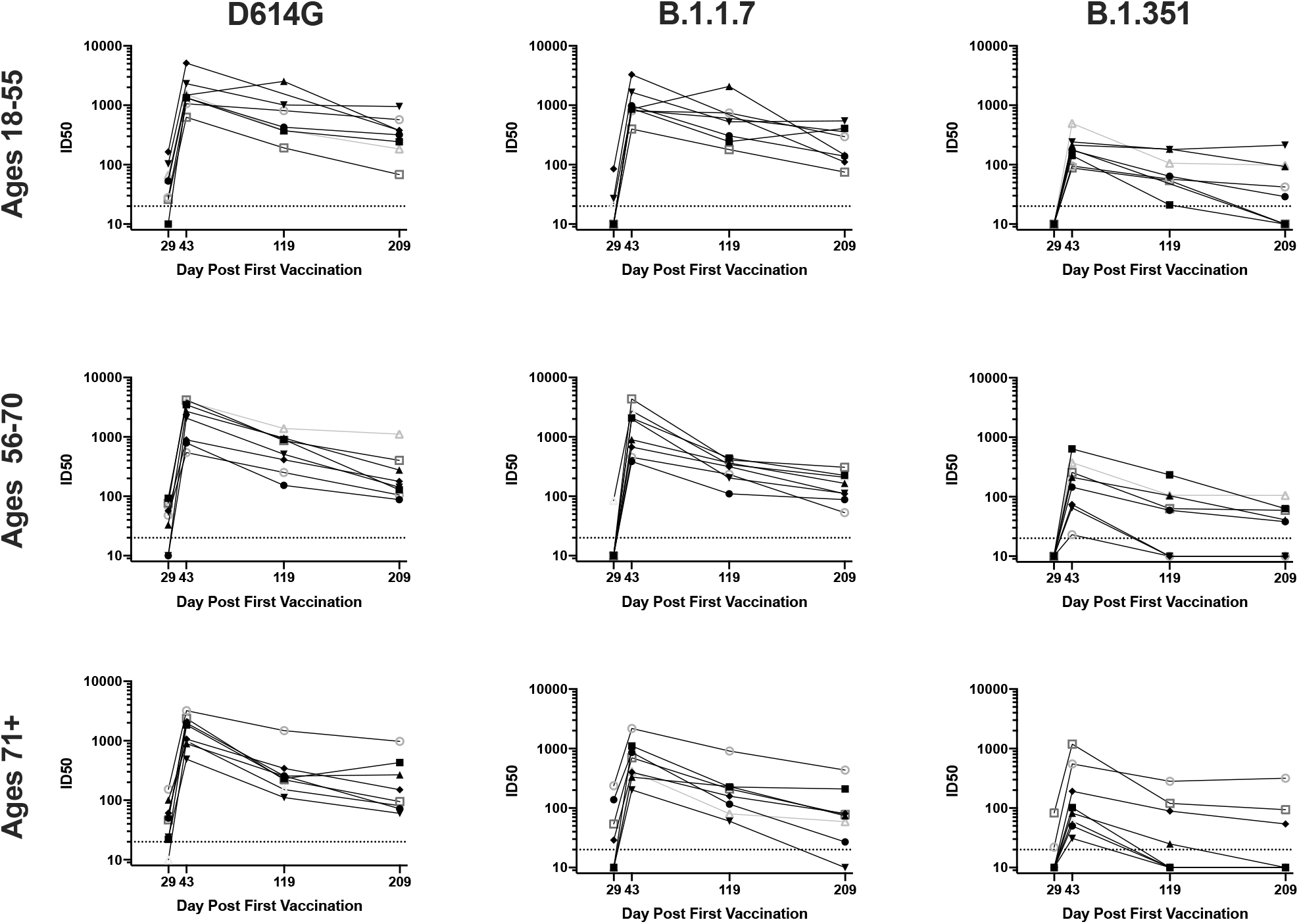
Neutralization of D614G and variant pseudoviruses is sustained for 6 months. 100 ug mRNA-1273 was delivered at Days 1 and 29. Each line represents the pseudovirus neutralization ID50s at Days 29 (4 weeks after first dose), 43 (two weeks after second dose), 119, and 209 for a single subject., n=8 per group. Neutralization activity of sera from subjects aged 18-55 (top row), 56-70 (middle row), and 71 + (bottom row) was measured against pseudoviruses bearing spike of D614G (left), B.1.1.7 (middle), or B.1.351 (right).

The experimental analyses were then expanded to additional variants and additional assays. Three functional assays and two binding assays were used to assess the humoral response to SARS-CoV-2 spike. SARS-CoV-2 neutralization was measured in two ways: the lentivirus-based pseudovirus assay, and a live-virus FRNT neutralization assay (*20*). The third functional assay was a novel MSD-ECLIA-based ACE2 blocking assay, which measured the ability of mRNA-1273 elicited antibodies present in sera to compete and inhibit binding of labelled soluble ACE2 to the specific RBD (WA1 or variant) spotted onto the MSD plate. In this assay, ACE2 blocking is dependent upon the ability of serum antibodies to compete the affinity of ACE2 for RBD. Spike-binding antibodies were measured using 2 different methodologies: binding to full-length, membrane-embedded spike on the surface of transfected cells followed by flow cytometry (*22*); and a novel MSD-ECLIA multiplex binding assay to simultaneously measure IgG binding against both the stabilized soluble spike protein S-2P (*23*) and RBD proteins derived from WA1 and the B.1.1.7, B.1.351, and P1 variants. All samples were assessed against either WA1 or D614G, and the B.1.1.7 and B.1.351 variants, in all of these orthogonal serology assays. In addition, all samples were tested for binding in the cell surface assay to P.1, B.1.429, and B.1.526, and by multiplex binding to P.1 proteins. A subset of samples was evaluated by pseudovirus neutralization against P.1, B.1.429, and B.1.526. The specific sequences used in each assay are defined in Supplemental Table 1.

Consistently across assays, low-level recognition of all variants was observed after a single dose in at least some individuals (Day 29). Robust activity against all variants was measured two weeks after the second dose (Day 43) with moderate declines over time through Day 209 (Figure 2). Titers were lower for all variants compared to WA1 or D614G, with the magnitude of the effect differing by assay. In comparing the assays, the most dramatic differences in responses by variant were noted in the ACE2 blocking assay, and the least in binding to S-2P (Supplemental table 2). While the scale of each assay was unique, the values obtained for each assay on a per-sample basis correlated well with each other (Supplemental fig 1).

**Figure 2.**
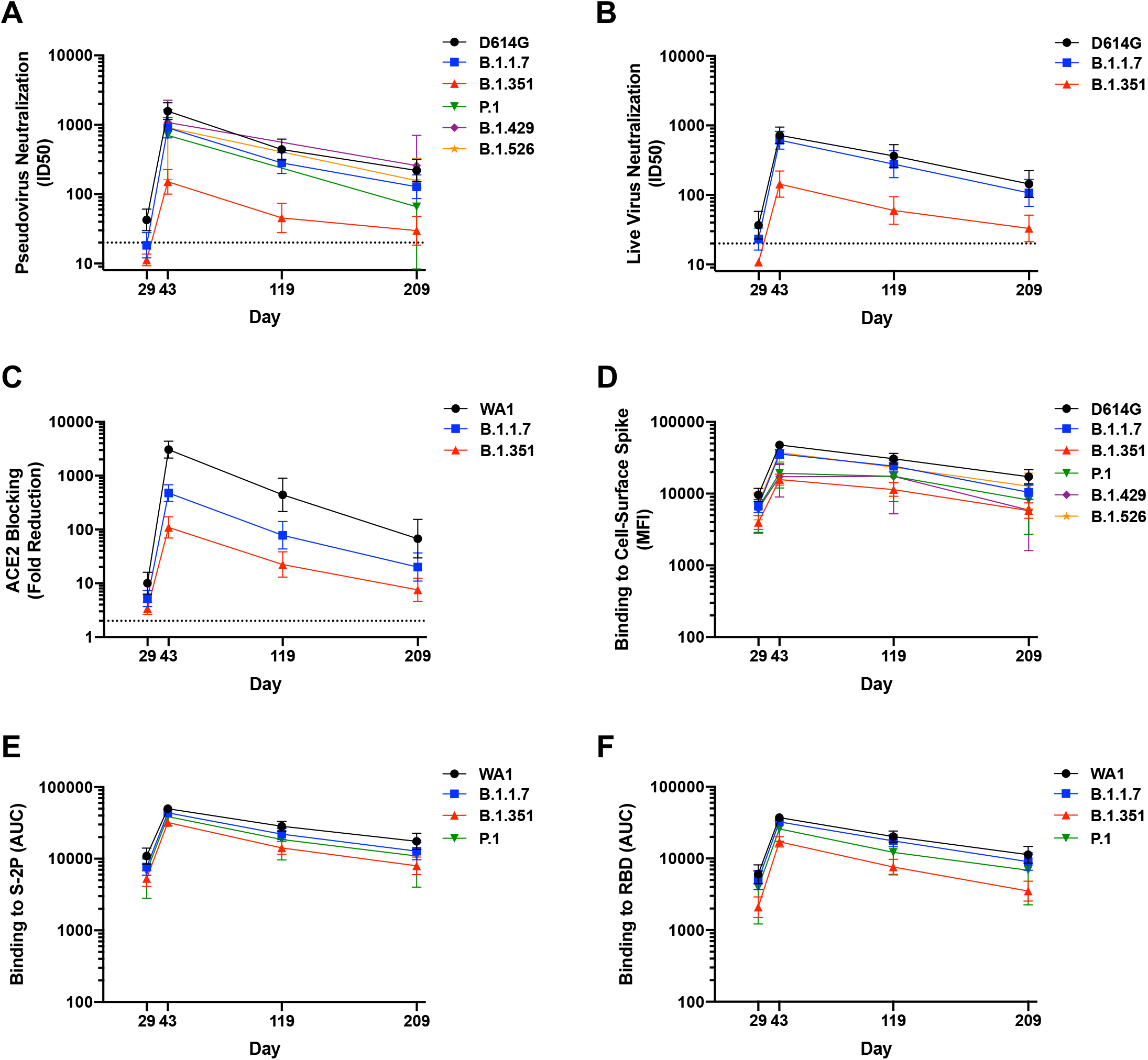
Binding and functional antibodies persist for 6 months following the second dose of mRNA-1273. For all assays, sera from n=24 individuals were sampled at 4 timepoints. Subjects were vaccinated with 100 ug mRNA-1273 at Days 1 and 29. **A**, Pseudovirus neutralization, expressed as 50% inhibitory dilution (ID50). Dotted line, limit of detection (>20). **B**. Live-virus FRNT neutralization, expressed as 50% inhibitory dilution (ID50). Dotted line, limit of detection (>20). **C**. Binding to cell-surface expressed full length spike, measured by flow cytometry and expressed as median fluorescence intensity (MFI). **D**. Serum blocking of ACE2 binding to RBD, measured by MSD-ECLIA and expressed as fold reduction of ACE2 binding. Dotted line, limit of detection (>2). **E**. Binding to soluble spike protein S-2P, measured by MSD-ECLIA and expressed as area under the curve (AUC). **F**. Bindng to receptor-binding domain protein (RBD), measured by MSD-ECLIA and expressed as area under the curve (AUC).

To quantify the breadth of responses, we calculated the number of sera that maintained detectable antibody titers in each assay and timepoint (Figure 3). Antibodies that bound to S-2P and RBD of WA1, B.1.1.7, B.1.351, and P.1 sequences were detected in all subjects at all timepoints. Likewise, binding to full-length cell-surface expressed spike was detected against D614G and all five variants at all timepoints. In contrast, the functional assays revealed deficits in antibody recognition of the variants. In the pseudovirus neutralization assay, 83% of Day 29 sera neutralized D614G, but 33% neutralized B.1.1.7 and only 8% could neutralize B.1.351. All Day 43 sera neutralized D614G and all five variants, however this cross-reactivity was reduced over time. All Day 209 sera neutralized D614G and B.1.429 in this assay, but fewer sera neutralized the other variants, with 96%, 88%, 85%, and 54% of sera neutralizing B.1.1.7, B.1.526, P.1, and B.1.351 respectively. Similarly, in the live virus assay, most Day 29 sera neutralized D614G and B.1.1.7 but only 8% neutralized B.1.351; all sera were active against all three at Day 43; and at Day 209, all sera neutralized D614G, 88% of sera neutralized B.1.1.7, and 58% neutralized B.1.351. The ACE2 blocking assay also showed reduced activity against B.1.351 at early and late timepoints (Figure 3). Thus, while all subjects showed activity against all variants in all assays two weeks after the second dose (Day 43), the functional assays revealed a decreased frequency of sera with detectable activity against B.1.351 and other variants after a single dose or 6 months after the second dose.

**Figure 3.**
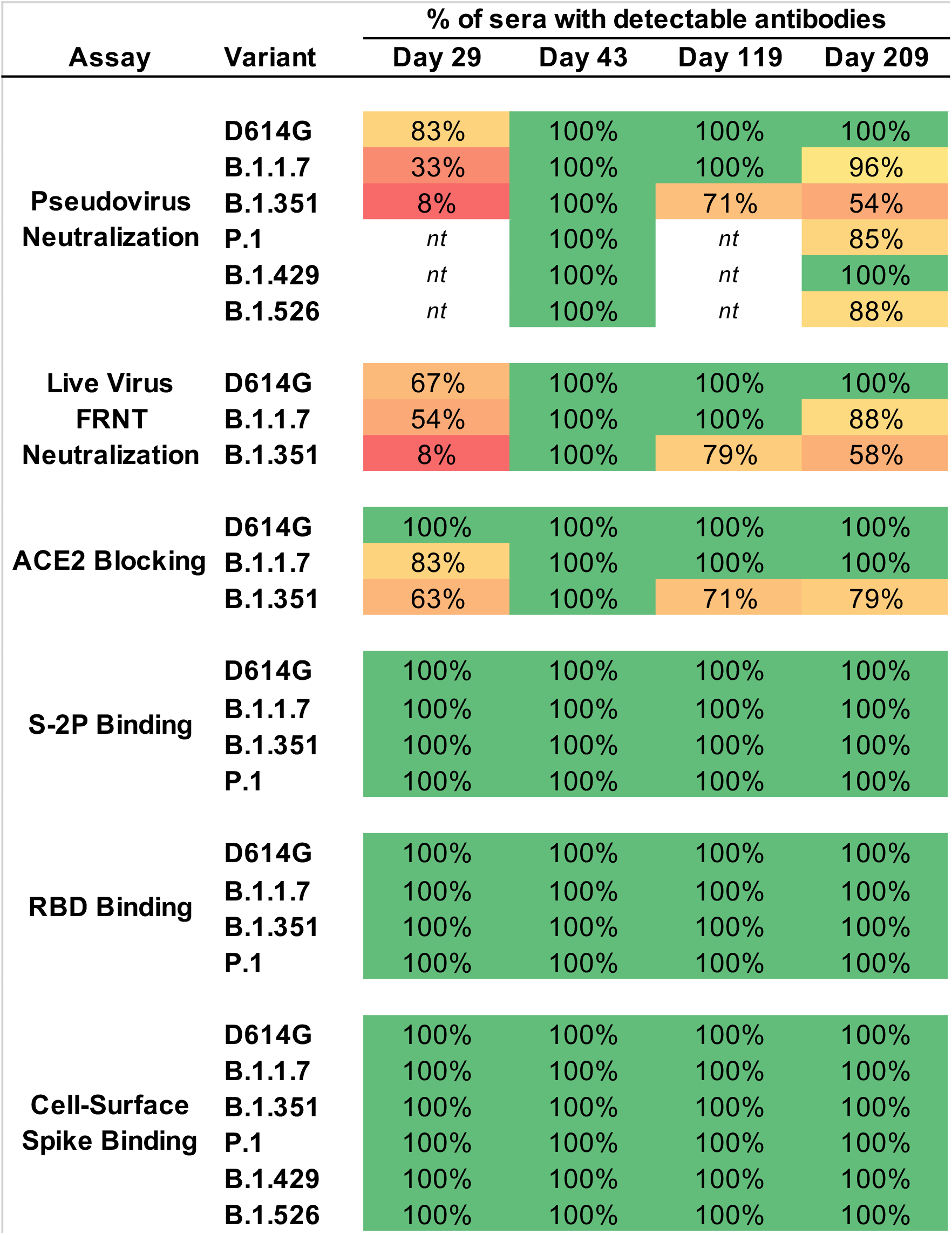
The majority of Day 209 sera maintain activity against WA1, D614G, and variants. Values are the percentage of sera (n=24 at each timepoint) for which antibodies were detected, for each variant. For pseudovirus and live-virus neutralization, samples were called detectable at ID50>20; for ACE2 blocking, at 2-fold decrease in signal; for S-2P and RBD binding, AUC>100; for cell-surface spike binding, MFI>100. *nt*, not tested.

In addition to the frequency of detectable responses, we quantified the magnitude of the change in titers against each variant at each timepoint. For pseudovirus neutralization, at timepoints after the second dose, the greatest differences were noted for B.1.351, which showed geometric mean decreases of 9.1-fold, 9.7-fold, and 7.0-fold at Days 43, 119, and 209 respectively (Figure 4) compared to D614G. A minimal effect of B.1.1.7 was observed in this assay, with less than 2-fold decrease at these timepoints. We assayed additional variants at Days 43 and 209, to capture the peak and 6-month timepoints. At Days 43 and 209, titers to B.1.429 decreased less than 2-fold; 2.1-fold and 2.4-fold-fold differences were noted for B.1.526, and 2.8- and 3.8-fold differences were measured for P.1. Live virus neutralization assays showed similar results: while the geometric mean titers against B.1.1.7 were less than 2-fold reduced compared to the D614G isolate at all timepoints, B.1.351 was 3.4-fold, 5.0-fold, 6.1-fold, and 4.3-fold less sensitive than D614G at the four timepoints respectively. In comparison, the ability of vaccine sera to prevent ACE2 binding to the SARS-CoV-2 spike RBD varied over a larger dynamic range. The loss of ability of mRNA-1273 vaccine sera to prevent recognition of ACE2 was greatest at Day 43: blocking of ACE2 binding to RBD of B.1.1.7 was 6.4-fold less than to RBD of WA1, and to RBD of B.1.351 was 28-fold less. As titers waned, the differences were smaller, with a 9.1-fold difference between RBDs of WA1 and B.1.351 at Day 209 (Figure 4). Thus, while the magnitude of the effects differed between these functional assays, the trends were the same, with mutations in B.1.1.7 spike region having minimal effect and B.1.351 showing a much greater impact; and P.1, B.526, and B.1.429 having an intermediate phenotype.

**Figure 4.**
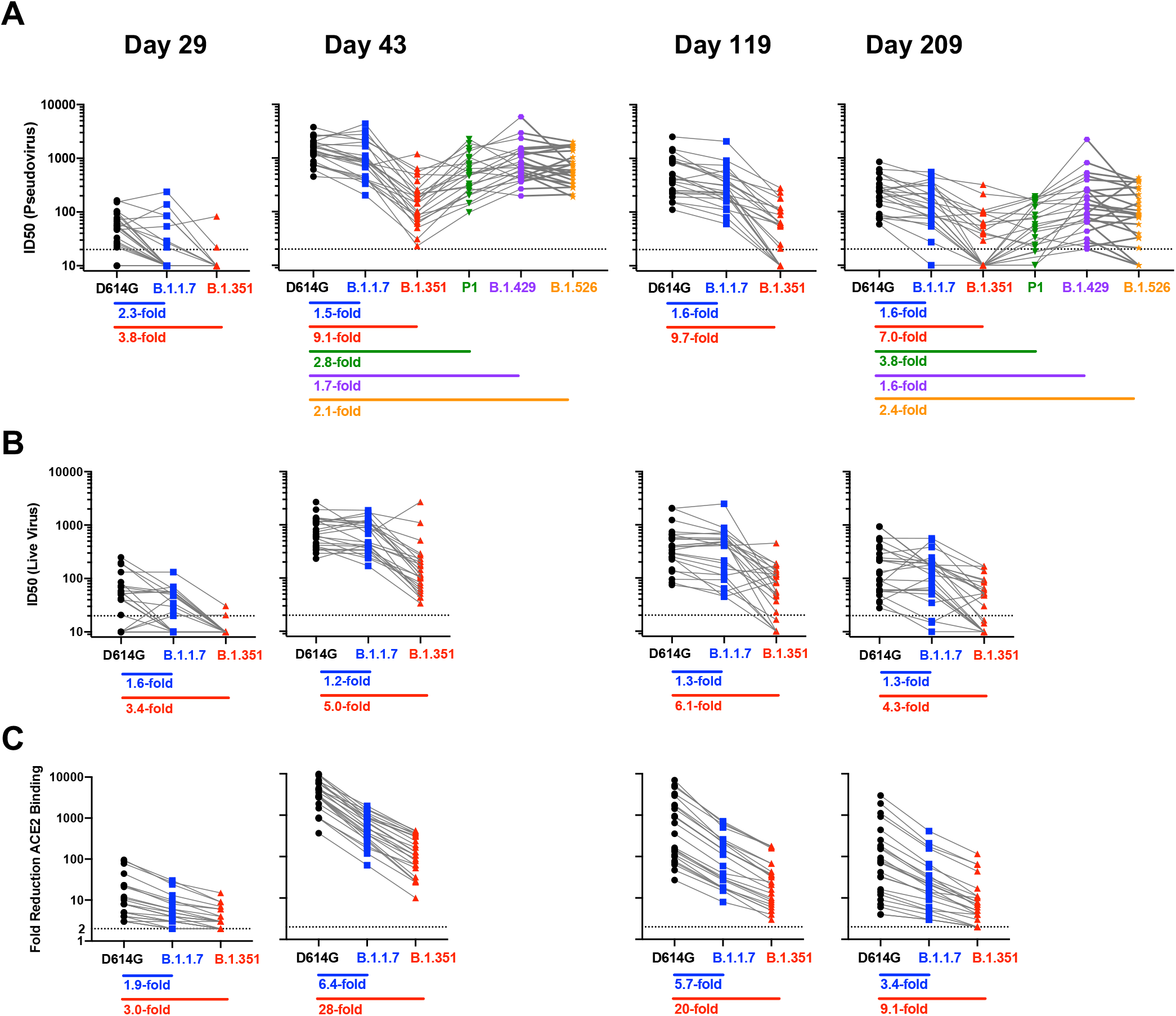
Impact of variants on antibody functions is stable over time. Sera from n=24 individuals were sampled at each timepoint. Fold change: geometric mean of ratios for each sample. **A**. ID50 in pseudovirus neutralization assays using D614G compared to B.1.1.7, B.1.351, P1, B.1.526, and B.1.429. **B**. ID50 in live virus FRNT neutralization assays using 83E (D614G) compared to B.1.1.7 or B.1.351. **C**. Blocking of ACE2 binding to WA1 RBD compared to B.1.1.7 RBD or B.1.351 RBD.

Similar trends were observed for the antibody binding assays, however the relative differences between variants were less pronounced (Figure 5). Binding to S-2P of B.1.1.7 and P.1 differed from binding to WA1 S-2P by 2.0-fold or less at all timepoints, while binding to S-2P of B.1.351 was no more than 2.2-fold diminished. The differences between variants were greater for binding to RBD (Figure 5), with 2.9-fold, 2.2-fold, 2.7-fold, and 3.2-fold decreased binding to RBD of B.1.351 compared to WA1 at Days 29, 43, 119, and 209 respectively. The antibody binding to RBD of the P.1 variant was minimally affected compared to the B.1.351 variant. Of note, the RBD of P.1 and B.1.351 differ by a single amino acid: B.1.351 has a K417N mutation relative to WA1, while the same position is K417T for P.1. The differences were similar for cell-surface binding, with binding to B.1.351 and B.1.426 reduced up to 3.0-fold compared to D614G.

**Figure 5.**
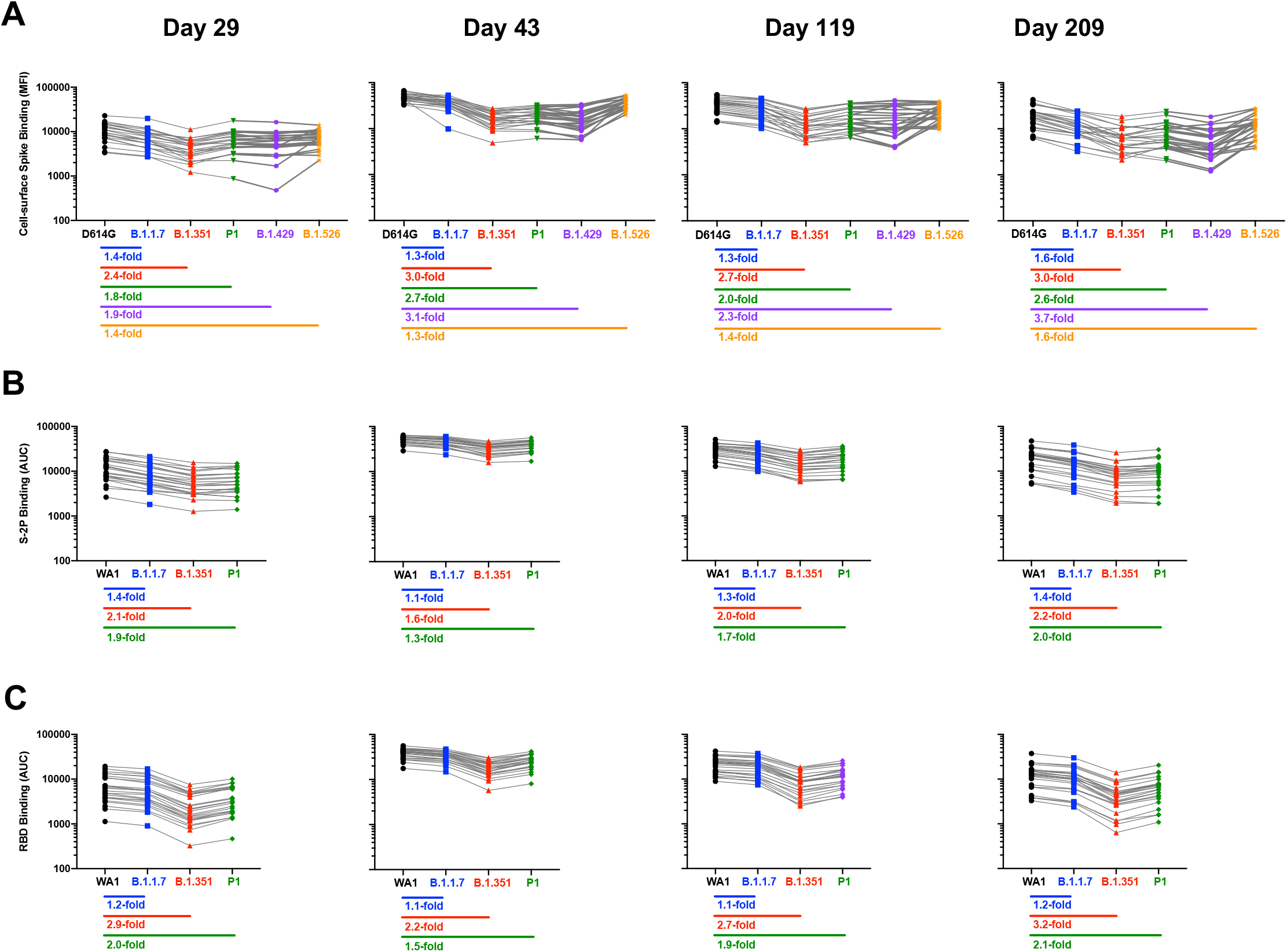
Impact of variants on antibody binding is stable over time. Sera from n=24 individuals were sampled at each timepoint. Fold change: median of ratios for each sample. **A**. Binding of cell-surface expressed full-length spike of D614G compared to variants. **B**. Binding to S-2P of WA1 compared to variants. **C**. Binding to RBD of WA1 compared to variants.

To understand the contributions of individual mutations to the immune escape noted in the variants of concern, we assayed Day 43 sera against pseudoviruses bearing D614G and one additional mutation: N501Y, which is present in both the B.1.1.7 and B.1.351 variants; N439K, previously shown to cause resistance to the therapeutic monoclonal antibody REGN 10987 (*24*), and Y453F, found in mink cluster 5 variants (*25*) and previously shown to cause resistance to the therapeutic monoclonal antibody REGN 10933 (*26*). None of these three showed a significant impact on neutralization by Day 43 sera (Fig S2). In contrast, E484K, which is present in B.1.351, P.1, and B.1.526, significantly impacted neutralization sensitivity, with a geometric mean 2.4-fold lower ID50. (Supplemental Figure 2).

Given importance of vaccines for older individuals, we examined whether the impact of variants differed between the age groups in this study. For each assay, we examined the fold-change in activity between each variant compared to WA1 or D614G for each sample, stratified by age. The results varied by assay and by variant (Supplemental Fig 3). For B.1.351, which showed the largest differences in all assays, the 18-55 groups showed a significantly greater fold-decrease in live-virus neutralization compared to the 56-70 and 71+ groups. However, there were no significant differences in pseudovirus neutralization between the groups, and ACE2 blocking ratios only differed between the 18-55 and 71+ groups. We previously reported a significant difference between the youngest and oldest groups in live-virus neutralization of WA1 at Day 209 (*18*). We therefore focused on that timepoint, and noted sporadic differences between age groups, but no overarching trends (Supplementary Figure 4). Overall, we did not observe any consistent differences between the age groups.

Across the various assays, the level of resistance of the variants showed a consistent hierarchy. As summarized in Figure 6, B.1.351 was the most resistant, B.1.1.7 was the least, and P.1, B.1.429, and B.1.526 had an intermediate phenotype in all assays. These data suggest that B.1.1.7 and B.1.351 bracket the magnitude of impacts of these variants.

**Figure 6.**
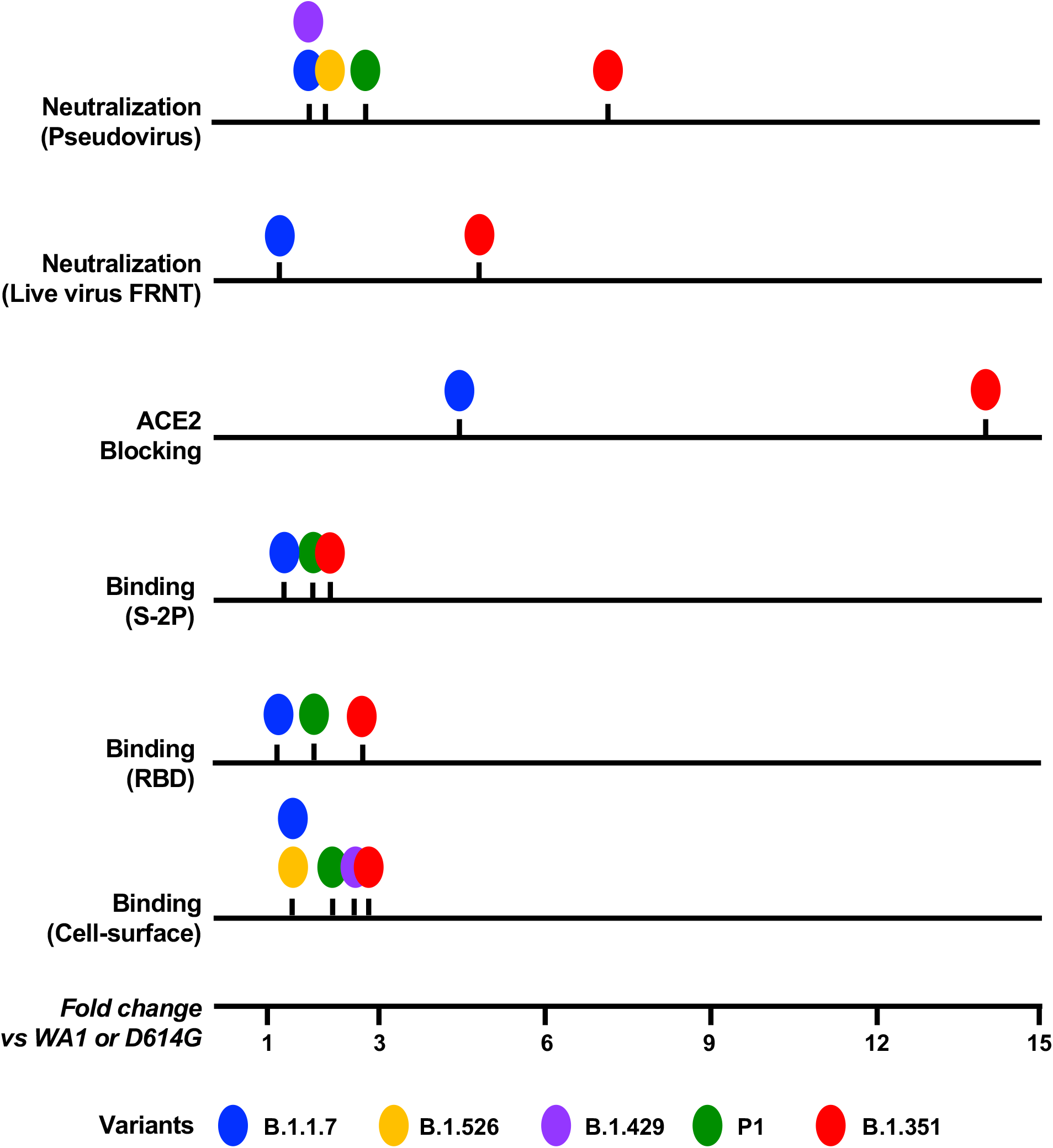
Rank-order of the impact of each variant is similar across assays. Individual lines and markers show that while the magnitude of the median fold change (compared to WA1/D614G) varies for individual assays, the rank order of impact on antibody recognition by these variants is similar across all assays tested. Median fold change was calculated for all samples. See also Supplemental Table 2.

## Discussion

SARS-CoV-2 Variants of Concern and Variants of Interest have been shown to escape from neutralizing antibodies (*3, 5-7, 10, 27-29*). Here, we present a comprehensive assessment of the impact of SARS-CoV-2 virus variants on antibody binding, ACE2 blocking, and neutralization. Over time, while the magnitude of the effects varied by methodology, the overall trend was consistent: mutations present in the B.1.1.7 variant had a minimal impact, while mutations in B.1.351 had a considerably greater effect upon antibody recognition and function, ranging from 3-to 15-fold depending on the assay. In addition, variants P.1, B.1.429, and B.1.526 showed an intermediate effect.

No single assay measurement, or antibody function, has been definitively identified as a correlate of protection from SARS-CoV-2 disease (*30, 31*). Many SARS-CoV-2 neutralization assays have been developed in response to the immediate requirements during the pandemic; varied live-virus and pseudovirus assays use different target cells, virus backbones, and methods of enumeration, they differ in sensitivity, and may even be measuring different modes of viral entry (for example, cell-membrane vs endosomal pathways). Measurements of antibody binding assays may be more sensitive than neutralization assays, with a larger dynamic range. Therefore, in this context, using diverse orthogonal assays in parallel can provide a better overall characterization of the humoral response. Here we show that results from binding to three forms of spike (cell surface, stabilized soluble S-2P, and RBD), while different in scale, show similar rank-order of antibody activity against the SARS-CoV-2 variants and similar dynamics over 7 months after the first vaccination. The same was true for two neutralization assays (pseudovirus and live virus) and ACE2 blocking. In the absence of a correlate of protection determined by a single methodology, data from multiple parallel assays will continue to be critically valuable for the assessment of CoVID-19 vaccines. Finally, we do not exclude the possibility that other functional characteristics of antibodies, not measured here, may contribute to protection.

The magnitude of the impact of these SARS-CoV-2 variants, and the mutations therein, differed with the format of the assay. The antibody binding differences between variants were more pronounced for RBD compared to the full-length soluble spike protein, S-2P, likely because the polyclonal response to the full length spike protein includes targeting of more epitopes, including conserved epitopes. Likewise, cell-surface expressed spike contains additional epitopes compared to S-2P, and is present in the membrane-bound context; in the assay, the differences between variants were slightly more pronounced than for S-2P binding. SARS-CoV-2 neutralizing activity, however, was more much affected by the variant sequences than was the polyclonal recognition of the spike protein itself, with many mRNA-1273 vaccine sera demonstrating loss of activity against B.1.351 in both neutralization assays. We speculate that the differential loss of neutralization compared to binding activity is due to immune pressure that favors mutations specifically at the epitopes against which neutralizing antibodies are elicited, for example E484K. Finally, ACE2 blocking was the assay that measured the greatest difference between variants. This methodology depends upon both the relative affinity of the variant RBD sequences for ACE2, and the affinity of the antibodies for binding to RBD at the same site as ACE2. The N501Y mutation, present in all of the tested variant RBD proteins, has been reported to increase affinity of RBD to ACE2 (*32, 33*). Thus, an antibody with the same binding affinity for WA1 and a variant RBD would be expected to have less capacity to block ACE2 binding to the variant; this will enhance the relative impact of variants on the blocking measurement. This assay also had the greatest dynamic range out of the methods used here, with upwards of 5000-fold blocking of ACE2 binding by some samples; consequently, despite large relative changes between variants, fewer sera showed complete loss of activity using this method.

Given the low availability of vaccine doses in many countries, there is interest in the immune responses after a single dose of vaccine. In this study, responses to the variants were limited after a single dose: at day 29 (4 weeks after the first dose), all subjects had binding antibodies against all variants tested, but only 2 of 24 sera (8%) could neutralize B.1.351 in pseudovirus or live-virus neutralization assays, and 33-54% could neutralize B.1.1.7 in the two assays respectively. While a single dose of mRNA-1273 provides partial protection against COVID-19 disease in the interval prior to the second vaccination (*34*), and similar data were reported for the mRNA vaccine BNT162b2 (*16, 17*), our observation of the limited magnitude and breadth of neutralizing activity at Day 29 underscores the importance of the full two-dose regimen of an mRNA vaccine for protection against SARS-CoV-2 variants.

Antibody activity against the most resistant variant, B.1.351, was substantially reduced over time – but not entirely abrogated. Indeed, at Day 209, more than half of sera (58%) still had detectable and robust neutralizing activity against B.1.351 in the pseudovirus assay, and 63% in the live-virus assay. All individuals retained binding activity; and 79% could block ACE2 binding to RBD of B.1.351. Importantly, all subjects had broadly cross-reactive activity against all variants at Day 43, the peak of the response. This indicates that the individuals in whom activity against the variants had waned to undetectable levels are likely to have memory B cells capable of responding to those variants in the event of exposure to virus or potentially with boosting vaccination, as seen in convalescent patients 6 months after infection (*35*).

Studies of SARS-CoV-2 specific antibody lineages, derived from memory B cells, sampled over time in COVID-19 convalescent individuals suggest that immune maturation months after infection can compensate for the variation seen in the SARS-CoV-2 genome (*35*). Moreover, several groups have reported an extremely robust response to vaccination in individuals who previously were infected with SARS-CoV-2, with notably increased neutralization titers against B.1.1.7 and B.1.351 (*11, 36, 37*). We therefore examined the relative recognition of variants at longitudinal timepoints after 2 doses of mRNA-1273 in COVID-19-naive individuals, however we did not see an increase in antibody breadth in this setting. The effects on antibody potency and breadth of a third dose of mRNA vaccine, encoding either the original (mRNA-1273) or the B.1.351 sequence (mRNA-1273.351) or co-administration of both, is currently under investigation: early results show strong boosting of responses to both D614G and variants by vaccination with either sequence (*38*).

Immune responses to vaccination are often weaker in older adults (*39*). In contrast, we previously showed that vaccination with mRNA-1273 elicited antibodies to SARS-CoV-2 WA1 in subjects aged 56 to 70 and 71 and older that are as potent (*20*) and durable (*18*) as those elicited in adults aged 18-55. Here we observed that responses to SARS-CoV-2 spike variants were statistically indistinguishable between age groups in most assays we, lending further support for the use of this vaccine to protect older populations. This supports the observed clinical data, where mRNA vaccines have strong protective effects against COVID-19 disease in the elderly (*17, 20*). However, some of the analyses performed here did show small differences, underscoring the importance of continued immune monitoring over time, particularly as new variants emerge, and the possibility that the aged populations may need more frequent boosting compared to younger individuals.

In summary, mRNA-1273-elicited neutralizing antibody activity against SARS-CoV-2 variants persisted six months after the second dose, albeit at reduced levels compared to WA1 and D614G, with more than half of subjects maintaining neutralizing activity against B.1.351 at the latest timepoint tested. High levels of binding antibodies recognizing B.1.351, as well as B.1.1.7, P.1, B.1.429, and B.1.526 were maintained in all subjects over this time period. The impact of variants on antibody recognition was consistent over time and across age groups. Additional studies will be needed to address the impact of new variants that will surely arise in areas of intense viral infection, such as B.1.617 variants (identified in India). While the correlates of vaccine-induced protection are not yet known, our data are encouraging for the use of this vaccine in the face of viral variation.

## Acknowledgements

We thank David Montefiori, Rosemarie Mason, Maryam Muhkamedova, Kathleen Neuzil, Cuiping Liu, and members of the Vaccine Research Center Virology Laboratory for helpful discussions. We thank Andy Pekosz for the B.1.351 variant virus, and Eli Boritz and Danny Douek for sequencing and analysis of the B.1.351 variant virus stock. We thank Huihui Mu and Michael Farzan for the ACE2-overexpressing 293T cells, and Adrian Creanga for Vero-TMPRSS2 cells.

## Funding

Emory Executive Vice President for Health Affairs Synergy Fund Award (MSS)

Pediatric Research Alliance Center for Childhood Infections and Vaccines and Children’s Healthcare of Atlanta (MSS)

Woodruff Health Sciences Center 2020 COVID-19 CURE Award (MSS);

National Institutes of Health grant UM1AI148373 (LJ)

National Institutes of Health grants UM1AI148576, UM1AI148684, and NIH P51OD011132 (EA, NGR)

National Institutes of Health grant HHSN272201500002C (JA)

Intramural Research Program of the Vaccine Research Center, NIAID, NIH (JRM, AMcD, BSG)

Coalition for Epidemic Preparedness Innovation (HB, BL)

## Author Contributions

Conceptualization: NDR, AMcD, SEO, AP, MSS

Laboratory Investigation: SEO, LL, CAT, SDS, SO, WS, LW, BF, ESY

Laboratory Supervision: KC, BSG, JRM, NDR, AMcD, AP, MR, JRM, MSS

Clinical Investigation: LJ, EA, NGR, ATW, JEL, JA, BL, HB, CJL, PCR, PAR, MM, JHB, CAR

Writing – original draft: NDR, SEO, AP, AMcD

Writing – review & editing: LJ, EA, JEL, PCR, BL, LW, JRM, MSS, CJL, CAR

## Data and materials availability

All data are available in the main text or the supplementary materials.

## Supplementary Materials

Materials and Methods

Supplementary Text: Members of the mRNA-1273 study group

Figs S1 to S5

Tables S1 and S2

## Materials and Methods

### Subjects and samples

Subjects in this manuscript participated in a phase 1, dose-escalation, open-label clinical trial of mRNA-1273, as previously reported (*18-21*). 8 subjects each were randomly chosen from participants from age cohorts 18-55, 56-70, and 71+ years of age who received two doses of 100 mcg mRNA-1273. The trial was conducted at Kaiser Permanente Washington Health Research Institute in Seattle, WA, the Emory University School of Medicine in Atlanta, GA, and the National Institute of Allergy and Infectious Diseases (NIAID) Vaccine Research Center (VRC) at the National Institutes of Health Clinical Center in Bethesda, MD. Enrolled adults were healthy and provided informed consent prior to any study procedures. Neither PCR nor serology for SARS-CoV-2 was utilized in screening.

### Spike sequences

The Spike sequences used in the assays are shown in Supplemental Table 1. The exact sequence of B.1.351 spike differed at amino acid 246 between the pseudovirus and live-virus neutralization assays. To address this difference, we compared both spike versions in the pseudovirus assay; overall, there was a 1.3-fold difference, which did not reach statistical significance (Wilcoxon matched-pairs signed rank test) (Supplementary Figure 4).

### Cells and Viruses

VeroE6 cells were obtained from ATCC (clone E6, ATCC, #CRL-1586) and cultured in complete DMEM medium consisting of 1x DMEM (VWR, #45000-304), 10% FBS, 25mM HEPES Buffer (Corning Cellgro), 2mM L-glutamine, 1mM sodium pyruvate, 1x Non-essential Amino Acids, and 1x antibiotics. VeroE6-TMPRSS2 cells were kindly provided by Drs. Barney Graham and Adrian Creanga (Vaccine Research Center, NIH, Bethesda, MD). EHC-083E (D614G SARS-CoV-2) and B.1.1.7 variants were previously described (*8, 9*). The B.1.351 variant was provided by Dr. Andy Pekosz (John Hopkins University, Baltimore, MD). Viruses were propagated in Vero-TMPRSS2 cells to generate viral stocks. B.1.351 stock was sequenced by Eli Boritz and Daniel Douek (Vaccine Research Center, NIH, Bethesda MD). Viral titers were determined by focus-forming assay on VeroE6 cells. Viral stocks were stored at -80°C until use. Compared to WA1, viral isolate 83E contains the D614G mutation in spike, and several additional mutations elsewhere in the genome.

### Pseudovirus neutralization

Neutralization activity against SARS-2-CoV was measured in a single-round-of-infection assay with pseudotyped virus particles (pseudoviruses) as previously described (*20*). To produce SARS-CoV-2 pseudoviruses, an expression plasmid bearing codon-optimized SARS-CoV-2 full-length S plasmid was co-transfected into HEK293T/17 cells (ATCC#CRL-11268) cells with packaging plasmid pCMVDR8.2, luciferase reporter plasmid pHR′CMV-Luc (*40*) and a TMPRSS2 plasmid (*41*). Spike sequences were: WA1, also called Wuhan-1, Genbank #: MN908947.3; and mutants made in the same plasmid, as in Table 1. Pseudoviruses were mixed with serial dilutions of sera or antibodies and then added to monolayers of ACE2-overexpressing 293T cells (gift of Michael Farzan and Huihui Mu), in triplicate. Three days post infection, cells were lysed, luciferase was activated with the Luciferase Assay System (Promega), and relative light units (RLU) were measured at 570 nm on a Spectramax L luminometer (Molecular Devices). After subtraction of background RLU (uninfected cells), % neutralization was calculated as 100x((virus only control)-(virus plus antibody))/(virus only control). Dose-response curves were generated with a 5-parameter nonlinear function, and titers reported as the serum dilution or antibody concentration required to achieve 50% (50% inhibitory dilution [ID50]) or 80% (80% inhibitory dilution [ID80]) neutralization. The input dilution of serum is 1:20, thus, 20 is the lower limit of quantification.

### Focus Reduction Neutralization Assay

FRNT assays were performed as previously described (*42*). Briefly, samples were diluted at 3-fold in 8 serial dilutions using DMEM (VWR, #45000-304) in duplicates with an initial dilution of 1:10 in a total volume of 60 μl. Serially diluted samples were incubated with an equal volume of SARS-CoV-2 (100-200 foci per well) at 37°C for 1 hour in a round-bottomed 96-well culture plate. The antibody-virus mixture was then added to Vero cells and incubated at 37°C for 1 hour. Post-incubation, the antibody-virus mixture was removed and 100 µl of prewarmed 0.85% methylcellulose (Sigma-Aldrich, #M0512-250G) overlay was added to each well. Plates were incubated at 37°C for 24 hours. After 24 hours, methylcellulose overlay was removed, and cells were washed three times with PBS. Cells were then fixed with 2% paraformaldehyde in PBS (Electron Microscopy Sciences) for 30 minutes. Following fixation, plates were washed twice with PBS and 100 µl of permeabilization buffer (0.1% BSA [VWR, #0332], Saponin [Sigma, 47036-250G-F] in PBS), was added to the fixed Vero cells for 20 minutes. Cells were incubated with an anti-SARS-CoV spike primary antibody directly conjugated to biotin (CR3022-biotin) for 1 hour at room temperature. Next, the cells were washed three times in PBS and avidin-HRP was added for 1 hour at room temperature followed by three washes in PBS. Foci were visualized using TrueBlue HRP substrate (KPL, # 5510-0050) and imaged on an ELISPOT reader (CTL). Antibody neutralization was quantified by counting the number of foci for each sample using the Viridot program (*43*). The neutralization titers were calculated as follows: 1 - (ratio of the mean number of foci in the presence of sera and foci at the highest dilution of respective sera sample). Each specimen was tested in duplicate. The FRNT-50 titers were interpolated using a 4-parameter nonlinear regression in GraphPad Prism 8.4.3. Samples that do not neutralize at the limit of detection at 50% are plotted at 10 and was used for geometric mean calculations.

The B.1.351 isolate used in the FRNT neutralization assay differs from the version used in the pseudovirus assay at a single amino acid, with the pseudovirus spike containing the R246I mutation. To test the potential impact of this mutation, we compared neutralization in the pseudovirus assay of B.1.351 and B.1.351v2, the latter matching the variant used in the FRNT assay (see Supplemental Table 1). For 13 sera tested, the ID50 values differed 1.3-fold, within the range of error for this assay (Supplemental Figure 5).

### 10-plex MSD-ECLIA

Multiplexed Plates (96 well) precoated with SARS-CoV-2 spike S-2P (WA1), SARS-CoV-2 RBD (WA1), SARS-CoV-2 spike S-2P (B.1.351), SARS-CoV-2 N Protein (WA1), SARS-CoV-2 spike S-2P (B.1.117), SARS-CoV-2 spike S-2P (P.1), SARS-CoV-2 RBD (B.1.351), SARS-CoV-2 RBD (B.1.117), SARS-CoV-2 RBD (P.1) and BSA are supplied by the manufacturer. On the day of the assay, the plate is blocked for 60 minutes with MSD Blocker A (5% BSA). The blocking solution is washed off and test samples are applied to the wells at 4 dilution (1:100, 1:500, 1:2500 and 1:10,000) unless otherwise specified and allowed to incubate with shaking for two hours. Plates are washed and Sulfo-tag labeled anti IgG antibody is applied to the wells and allowed to associate with complexed coated antigen – sample antibody within the assay wells. Plates are washed to remove unbound detection antibody. A read solution containing ECL substrate is applied to the wells, and the plate is entered into the MSD Sector instrument. A current is applied to the plate and areas of well surface where sample antibody has complexed with coated antigen and labeled reporter will emit light in the presence of the ECL substrate. The MSD Sector instrument quantitates the amount of light emitted and reports this ECL unit response as a result for each sample and standard of the plate. Magnitude of ECL response is directly proportional to the extent of binding antibody in the test article. All calculations are performed within Excel and the GraphPad Prism software, version 7.0. Readouts are provided as Area Under Curve (AUC).

### ACE2 blocking assay

Multiplexed Plates (96 well) precoated with RBD from WA1, B.1.351 and B.1.1.7 SARS-CoV-2 antigen are supplied by the manufacturer. On the day of the assay, the plate is blocked for 30 minutes with MSD Blocker A (5% BSA). The blocking solution is washed off and test samples are applied to the wells at 1:10, 1:20 and 1:40 dilution unless otherwise specified and allowed to incubate with shaking for one hour. Sulfo-tag labeled ACE2 is applied to the wells and allowed to associate with sample and RBD within the assay wells. Plates are washed to remove unbound detection antibody. A read solution containing ECL substrate is applied to the wells, and the plate is entered into the MSD Sector instrument. A current is applied to the plate and areas of well surface where RBD has complexed with ACE2-SulfoTag will emit light in the presence of the ECL substrate. The MSD Sector instrument quantitates the amount of light emitted and reports this ECL unit response as a result for each sample and standard of the plate. The amount of signal emitted in wells containing no sample (assay diluent only) is evaluated as the maximal binding response. Reduction of ECL response from this maximal readout is directly proportional to the extent of competitive binding activity in the test article. All calculations are performed within Excel and the GraphPad Prism software, version 7.0. Each fold reduction readout is generated against the maximal signal for the matched RBD antigen.

### Cell-surface spike binding

HEK293T cells were transiently transfected with plasmids encoding full length SARS-CoV-2 spike variants using lipofectamine 3000 (L3000-001, ThermoFisher) following manufacturer’s protocol. After 40 hours, the cells were harvested and incubated with serum diluted 1:160 in PBS for 30 minutes. After incubation, the cells were washed and incubated with an allophycocyanin conjugated anti-human IgG (709-136-149, Jackson Immunoresearch Laboratories) along with a live/dead fixable aqua dead cell stain kit (ThermoFisher) for another 30 minutes. The cells were then washed and fixed with 1% paraformaldehyde (15712-S, Electron Microscopy Sciences). The samples were then acquired on a BD LSR Fortessa X-50 flow cytometer (BD biosciences) and analyzed using Flowjo (BD biosciences). For each serum sample, the median fluorescence intensity (MFI) in the allophycocyanin fluorescence channel was determined for only the spike-transfected cells (typically, 75-90% of all cells).

## Supplementary Text

### mRNA-1273 Study Group

The following study group members were all closely involved with the design, implementation, and oversight of the mRNA-1273 clinical trial.

<u>Division of Microbiology and Infectious Diseases, National Institute of Allergy and Infectious Diseases, National Institutes of Health, Bethesda, MD.</u> Jae Arega, M.S., John H. Beigel, M.D., Wendy Buchanan, M.S., B.S.N., Mohammed Elsafy, M.D., Binh Hoang, Pharm.D., Rebecca Lampley, M.Sc., Aparna Kolhekar, Ph.D., Hyung Koo, B.S.N., Catherine Luke, Ph.D., Mamodikoe Makhene, M.D., M.P.H., Seema Nayak, M.D., Rhonda Pikaart-Tautges, B.S., Paul C. Roberts, Ph.D., Janie Russell, B.S., Elisa Sindall, B.S.N.

<u>The Emmes Company, LLC, Rockville, MD</u>. Jim Albert, M.S., Pratap Kunwar, M.S., Mat Makowski, Ph.D.

<u>Emory University School of Medicine, Atlanta, GA</u>. Evan J. Anderson, M.D., Amer Bechnak, M.D., Mary Bower, R.N., Andres F. Camacho-Gonzalez, M.D., M.Sc., Matthew Collins, M.D., Ph.D., Ana Drobeniuc, M.P.H., Venkata Viswanadh Edara, Ph.D., Srilatha Edupuganti, M.D., M.P.H, Katharine Floyd, Theda Gibson, M.S., Cassie M. Grimsley Ackerley, M.D., Brandi Johnson, Satoshi Kamidani, M.D., Carol Kao, M.D.; Colleen Kelley, M.D., M.P.H., Lilin Lai, M.D., Hollie Macenczak, R.N., Michele Paine McCullough, M.P.H., Etza Peters, R.N., Varun K. Phadke, M.D., Paulina A. Rebolledo, M.D. M.Sc., Christina A. Rostad, M.D., Nadine Rouphael, M.D., Erin Scherer Ph.D., D.Phil., Amy Sherman, M.D., Kathy Stephens, R.N., Mehul S. Suthar, Ph.D., Mehgan Teherani, M.D., M.S., Jessica Traenkner, P.A., Juton Winston, Inci Yildirim, M.D., Ph.D.

<u>Kaiser Permanente Washington Health Research Institute, Seattle, WA.</u> Lee Barr, R.N., Joyce Benoit, R.N., Heather Beseler, M.B.A., Rachael Burganowski, M.S., Barbara Carste, M.P.H., Joe Choe, B.S., John Dunn, M.D., M.P.H., Maya Dunstan, M.S., R.N., Roxanne Erolin, M.P.H., Jana ffitch, L.P.N., Colin Fields, M.D., Lisa A. Jackson, M.D., Erika Kiniry, M.P.H., De Vona Lang, L.M.P., Susan Lasicka, R.Ph., Stella Lee, B.A., Matthew Nguyen, M.P.H., Jennifer Nielsen, M.N., A.R.N.P., Hallie Phillips, M.ed., Stephanie Pimienta, B.S., David Skatula, R.Ph., Janice Suyehira, M.D., Karen Wilkinson, M.N., A.R.N.P., Michael Witte, Pharm.D.

<u>Moderna, Inc., Cambridge, MA.</u> Hamilton Bennett, M.Sc., Nedim Emil Altaras, Ph.D., Andrea Carfi, Ph.D., Marjorie Hurley, Pharm.D., Brett Leav, M.D., Rolando Pajon, Ph.D., Wellington Sun, M.D., Tal Zaks, M.D., Ph.D.

<u>Seattle Children’s Research Institute, Seattle, WA.</u> Rhea N. Coler, M.Sc., Ph.D., Sasha E. Larsen, Ph.D.

<u>University of Maryland School of Medicine, Baltimore, MD.</u> Kathleen M. Neuzil, M.D.

<u>University of North Carolina, Durham, NC.</u> Lisa C. Lindesmith, M.S., David R. Martinez, Ph.D., Jennifer Munt, B.S., Michael Mallory, M.P.H., Caitlin Edwards, B.S., Ralph S. Baric, Ph.D.

<u>Vaccine Research Center, National Institute of Allergy and Infectious Diseases, National Institutes of Health, Bethesda, M.D.</u> Nina M. Berkowitz, M.P.H., Kevin Carlton, M.S., Kizzmekia S. Corbett, Ph.D., Pamela Costner, R.N., B.S.N., Nicole A. Doria-Rose, Ph.D., Britta Flach, Ph.D., Martin Gaudinski, M.D., Ingelise Gordon, R.N., Barney S. Graham, M.D., LaSonji Holman, F.N.P., Julie E. Ledgerwood, D.O., Kwanyee Leung, Ph.D., Bob C. Lin, B.S., Mark K. Louder, John R. Mascola, M.D., Adrian B. McDermott, Ph.D., Kaitlyn M. Morabito, Ph.D., Laura Novik, R.N., M.A., Sarah O’Connell, M.S., Sijy O’Dell, M.S., Marcelino Padilla, B.S., Amarendra Pegu, Ph.D, Stephen D. Schmidt, B.S., Phillip A. Swanson II, Ph.D., Chloe A. Talana, B.S., Lingshu Wang, Ph.D., Alicia T. Widge, M.D., M.S., Eun Sung Yang M.S., Yi Zhang B.S.

<u>Vanderbilt University Medical Center, Nashville, TN.</u> James D. Chappell, M.D., Ph.D., Mark R. Denison, M.D., Tia Hughes, M.S., Xiaotao Lu, M.S., Andrea J. Pruijssers, Ph.D., Laura J. Stevens, M.S.

<u>Fred Hutchinson Cancer Research Center, Seattle WA.</u> Christine M. Posavad, Ph.D

<u>University of Washington, Seattle, WA.</u> Michael Gale, Jr., Ph.D.

<u>University of Texas Medical Branch, Galveston, TX.</u> Vineet Menachery, Ph.D., Pei-Yong Shi, Ph.D.

**Fig. S1.**
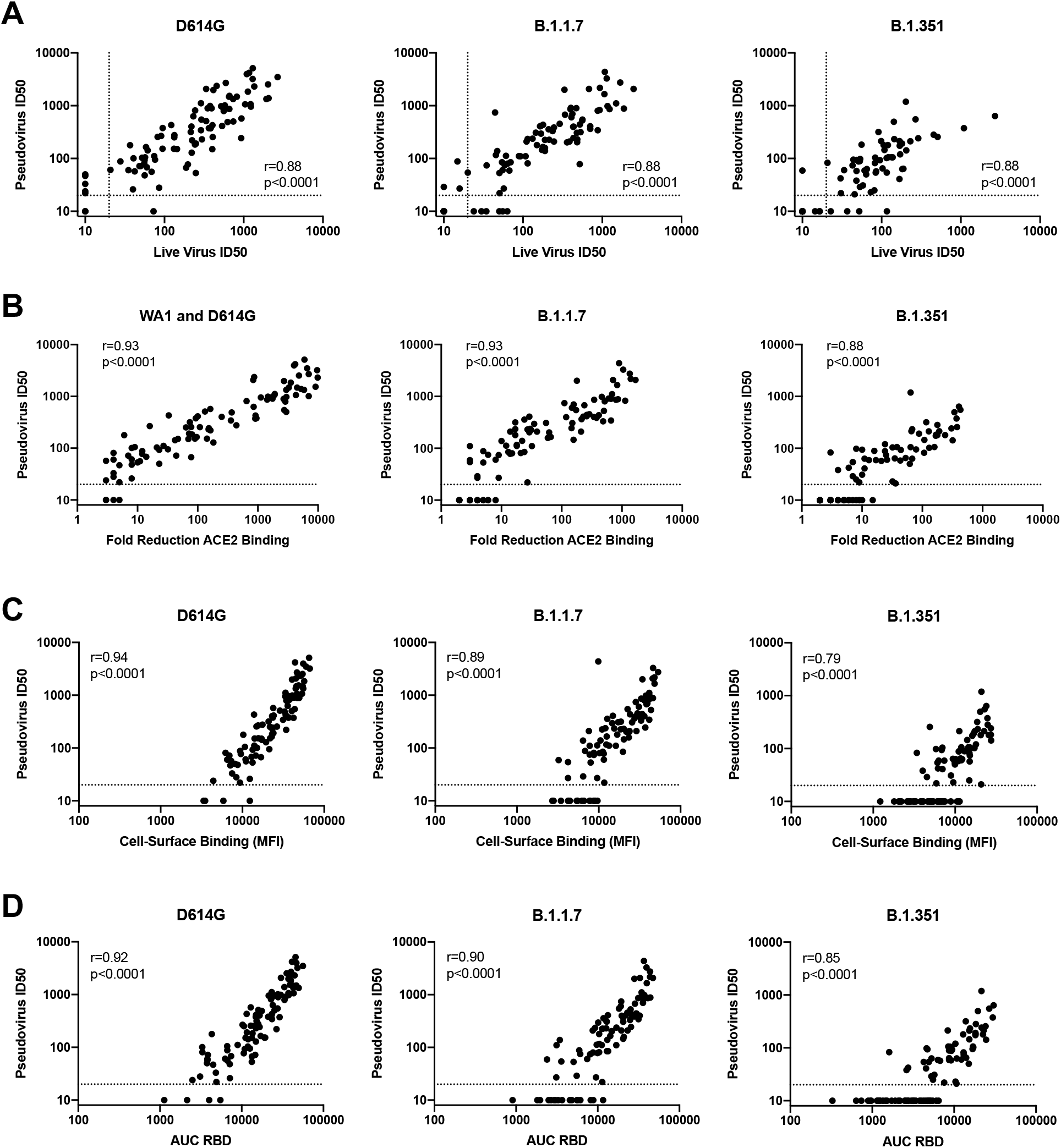
Functional and binding assays correlate well with each other. Each graph shows n=96 serum samples. r values: Spearman’s rho. Graphs show pseudovirus neutralization compared to: **A**, live-virus FRNT ID50, **B**. fold reduction in ACE2 binding, **C**. cell-surface binding median fluorescence intensity (MFI), **D**. binding to RBD in MSD-ECLIA assay, expressed as area under the curve (AUC). Left: WA1 or D614G; middle, B.1.1.7; right, B.1.351.

**Fig. S2.**
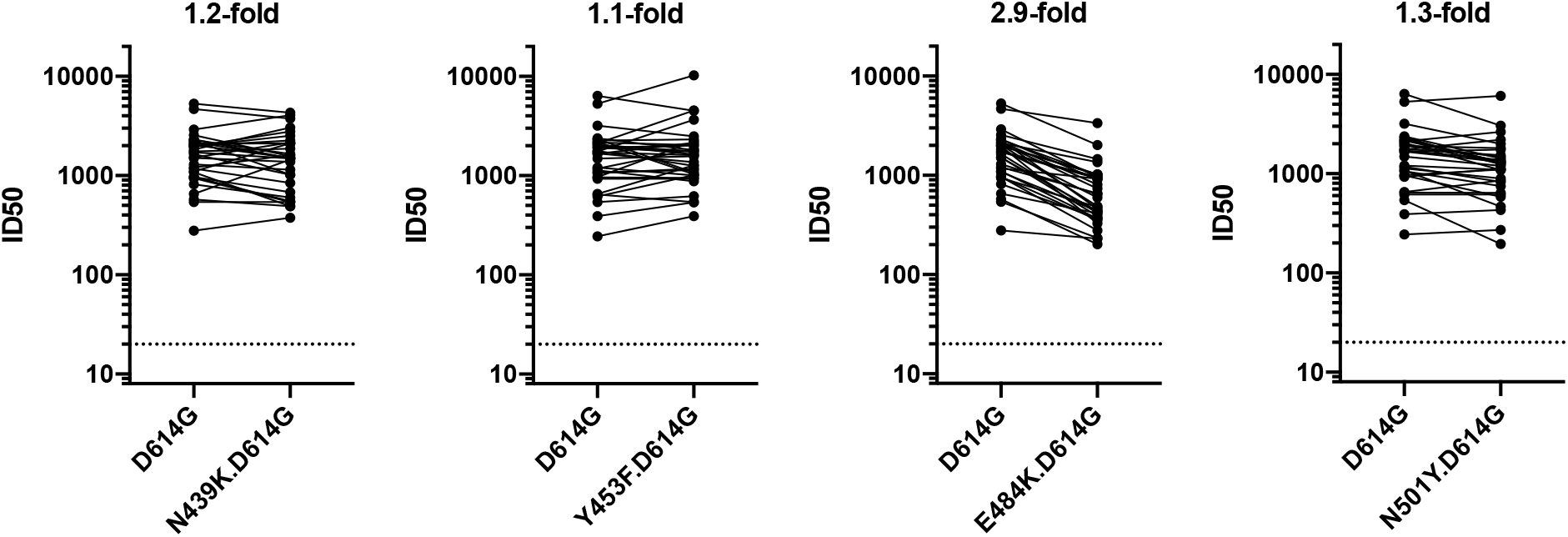
Point mutations cause modest decreases in neutralizing activity. Day 43 Sera were assessed in lentivirus-based pseudovirus neutralization assay. 33 sera were tested, inclusive of the 24 used in other figures plus additional samples as described in (20). Pseudoviruses were: D614G, D614G.N439K, D614G.Y453F, D614G.E484K, and D614G.N501Y. For each pair of viruses, the fold-differenec is the geometric mean of the ratio of ID50s for each serum.

**Fig. S3.**
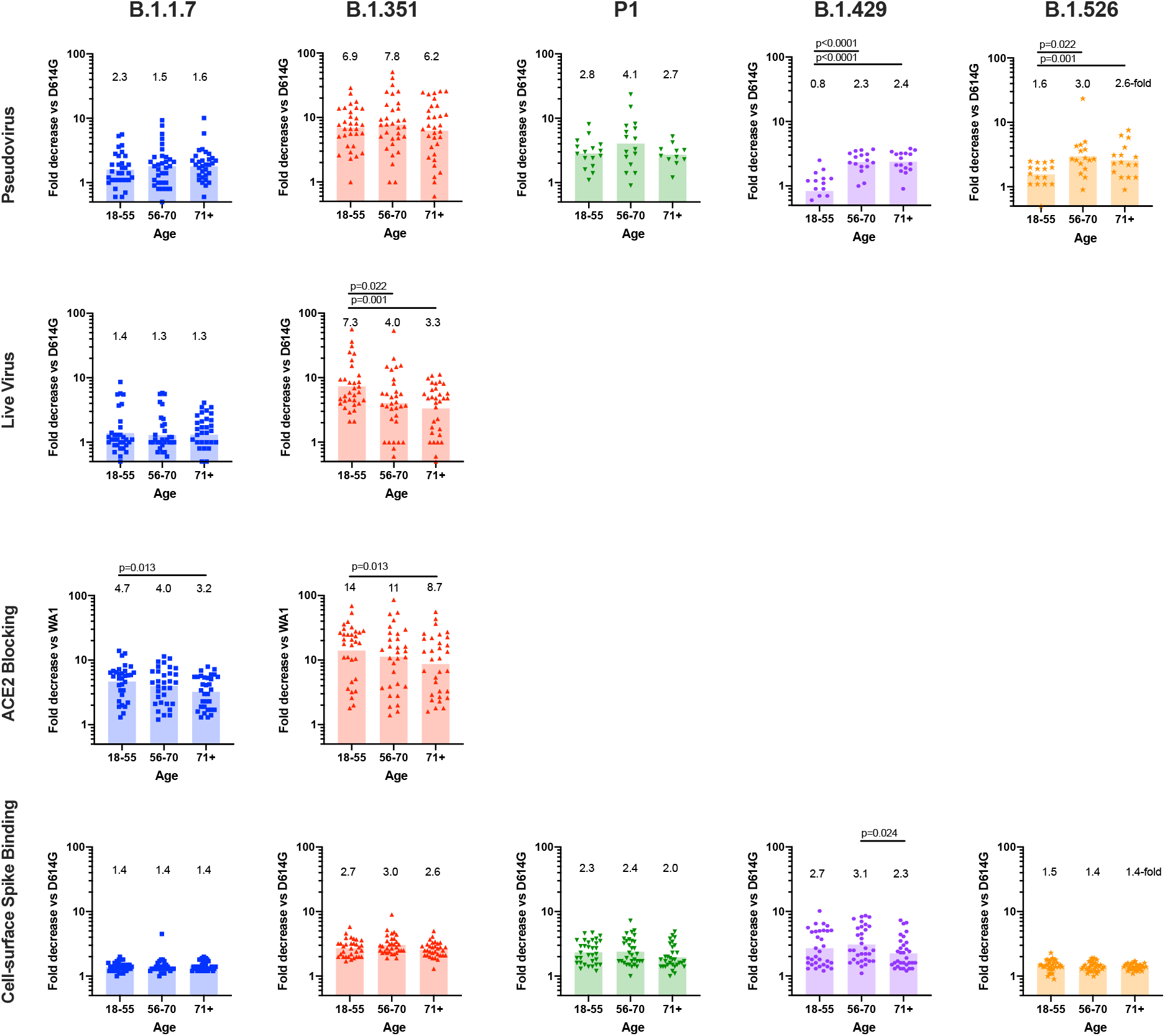
Effect of age on relative recognition of variants. Fold reduction in ID50 in each age group (8 subjects, 4 timepoints each, n=32 total) for each variant compared to WA1 or D614G. Bar: geometric mean. p values: Mann-Whitney test; values are not corrected for multiple comparisons; * p=0.01-0.05, ** p=0.001-0.01.

**Fig. S4.**
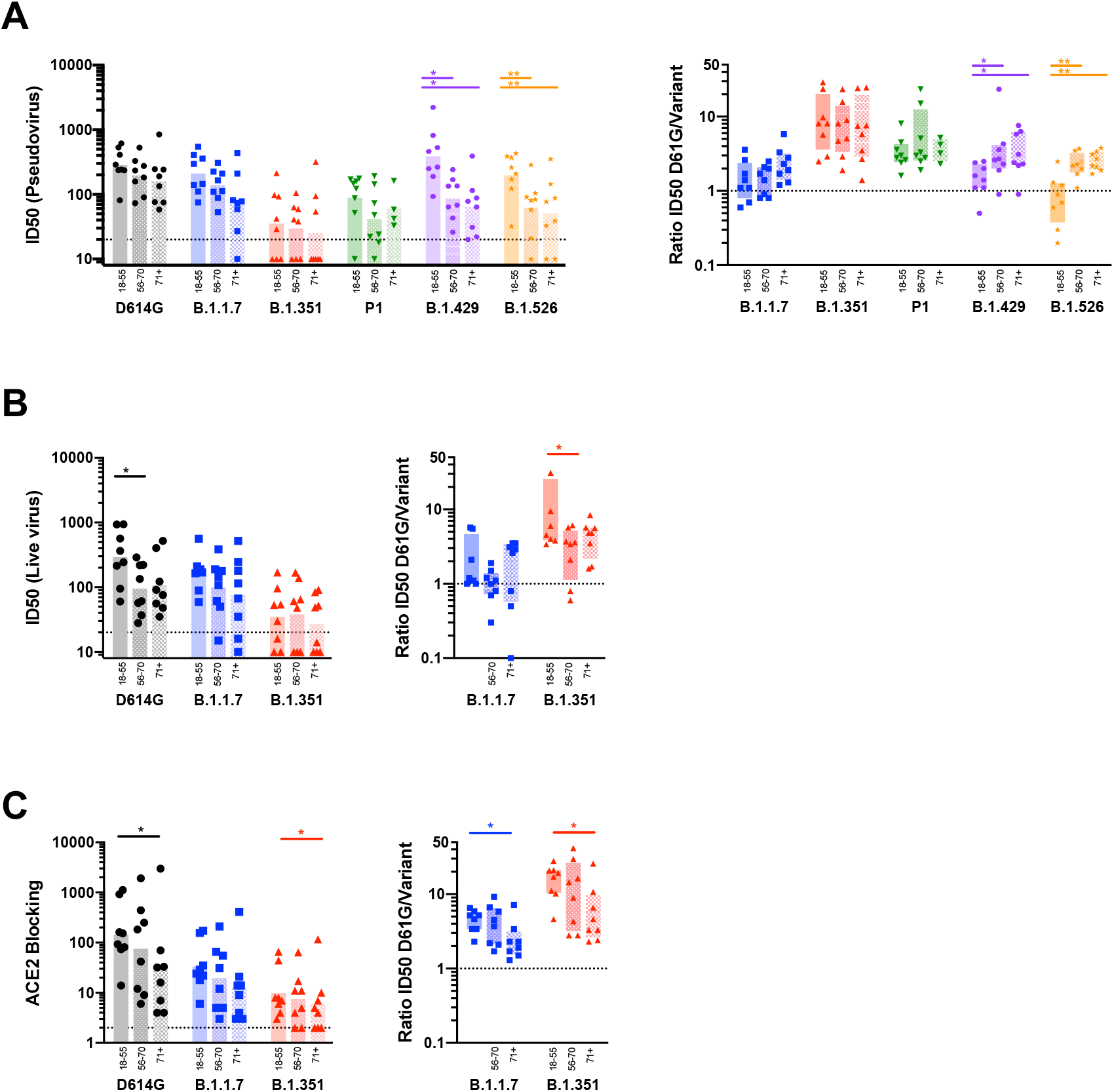
Effect of age on relative recognition of variants at Day 209. Left, assay values for each variant. Bar: geometric mean. Right, ratios compared to WA1 or D614G for each age group (n=8) and variant. p values: Mann-Whitney test; values are not corrected for multiple comparisons; * p=0.01-0.05, ** p=0.001-0.01. **A**. Pseudovirus neutralization. **B**. Live-virus FRNT neutralization. **C**. ACE2 blocking assay.

**Fig. S5.**
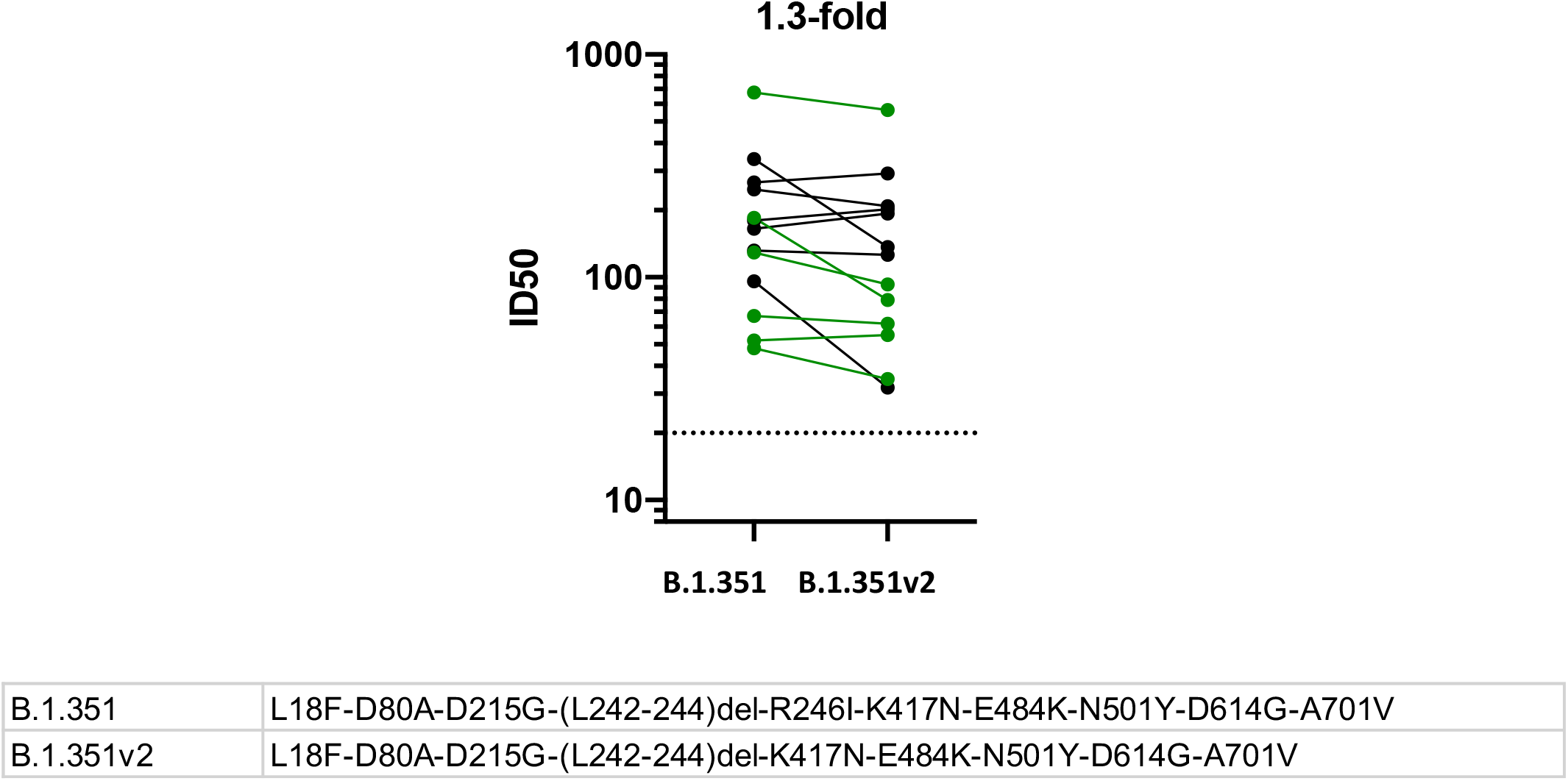
Two versions of B.1.351 yield similar pseudovirus neutralization IC50s. N=13 Sera (black, ages 18-55; green, ages 55-70) were assessed in pseudovirus neutralization assay. The spike proteins in the pseudoviruses differ only at amino acid 246 as indicated.

**Table S1.** Sequences of spike proteins used in each assay.

|  |  |
| --- | --- |
| B.1.351 | L18F-D80A-D215G-(L242-244)del-R246I-K417N-E484K-N501Y-D614G-A701V |
| B.1.351v2 | L18F-D80A-D215G-(L242-244)del-K417N-E484K-N501Y-D614G-A701V |

**Table S1.**
| **Pseudovirus** | **Name** | **Mutations relative to parental WA1** |
| --- | --- | --- |
|  | D614G | D614G |
|  | D614G.N439K | N439K-D614G |
|  | D614G.Y453F | Y453F-D614G |
|  | D614G.E484K | E484K-D614G |
|  | D614G.N501Y | N501Y-D614G |
|  | B.1.1.7 | d69H/70V-Y144del-N501Y-A570D-D614G-P681H-T716I-S982A-D1118H |
|  | B.1.351 | L18F-D80A-D215G-(L242-244)del-R246I-K417N-E484K-N501Y-D614G-A701V |
|  | B.1.351v2 | L18F-D80A-D215G-(L242-244)del-K417N-E484K-N501Y-D614G-A701V |
|  | P.1 | L18F-T20N-P26S-D138Y-R190S-K417T-E484K-N501Y-D614G-H655Y-T1027I-V1176F |
|  | B.1.429 | S13I-W152C-L425R-D614G |
|  | B.1.526 | L5F-T95I-D253G-E484K-D614G-A701V |
| **Cell-surface expressed Spike** | D614G | D614G |
|  | B.1.1.7 | d69H/70V-Y144del-N501Y-A570D-D614G-P681H-T716I-S982A-D1118H |
|  | B.1.351 | L18F-D80A-D215G-(L242-244)del-R246I-K417N-E484K-N501Y-D614G-A701V |
|  | P.1 | L18F-T20N-P26S-D138Y-R190S-K417T-E484K-N501Y-D614G-H655Y-T1027I-V1176F |
|  | B.1.429 | S13I-W152C-L425R-D614G |
|  | B.1.526 | L5F-T95I-D253G-E484K-D614G-A701V |
| **S-2P Proteins** | WA1 | D614G |
|  | B.1.1.7 | d69H/70V-Y144del-N501Y-A570D-D614G-P681H-T716I-S982A-D1118H |
|  | B.1.351 | L18F-D80A-D215G-(L242-244)del-R246I-K417N-E484K-N501Y-D614G-A701V |
|  | P.1 | L18F-T20N-P26S-D138Y-R190S-K417T-E484K-N501Y-D614G-H655Y-T1027I-V1176F |
| **RBD Proteins** | WA1 |  |
|  | B.1.1.7 | N501Y |
|  | B.1.351 | K417N-E484K-N501Y |
|  | P.1 | K417T-E484K-N501Y |
| **Live Virus** | 83E (D614G)<sup>1</sup> | D614G |
|  | B.1.1.7 | d69H/70V-Y144del-N501Y-A570D-D614G-P681H-T716I-S982A-D1118H |
|  | B.1.351<sup>2</sup> | L18F-D80A-D215G-(L242-244)-K417N-E484K-N501Y-D614G-A701V |

<sup>1</sup>Live virus strain 83E differs from WA1 at a single position in Spike, D614G; there are other differences across the genome, as reported in (Edara et al, 2021 JAMA)
<sup>2</sup>Live virus strain B.1.351 has same spike mutations as B.1.351v2 used for pseudoviruses.

**Table S2.** Geometric mean of ratios of value for D614G (pseudovirus, live virus, cell-surface spike binding) or WA1 (ACE2 blocking, S-2P binding, RBD binding) compared to the indicated variant.

| Assay | Variant | Ratio |
| --- | --- | --- |
| Pseudovirus<br>Neutralization | B.1.1.7 | 1.7 |
|  | B.1.351 | 6.9 |
|  | P.1 | 3.2 |
|  | B.1.429 | 1.7 |
|  | B.1.526 | 2.3 |
| Live Virus<br>Neutralization | B.1.1.7 | 1.3 |
|  | B.1.351 | 4.6 |
| ACE2 Blocking | B.1.1.7 | 3.9 |
|  | B.1.351 | 11.1 |
| S-2P Binding | B.1.1.7 | 1.3 |
|  | B.1.351 | 1.9 |
|  | P.1 | 1.7 |
| RBD Binding | B.1.1.7 | 1.2 |
|  | B.1.351 | 2.7 |
|  | P.1 | 1.9 |
| Cell-Surface<br>Spike Binding | B.1.1.7 | 1.4 |
|  | B.1.351 | 2.8 |
|  | P.1 | 2.2 |
|  | B.1.429 | 2.6 |
|  | B.1.526 | 1.4 |

